# Quantification of membrane binding and diffusion using Fluorescence Correlation Spectroscopy diffusion laws

**DOI:** 10.1101/2022.09.12.507540

**Authors:** Anita Mouttou, Erwan Bremaud, Julien Noero, Rayane Dibsy, Coline Arone, Johnson Mak, Delphine Muriaux, Hugues Berry, Cyril Favard

**Author notes:** Equivalent Contribution: *Anita Mouttou, Erwan Bremaud.

## Abstract

Many transient processes in cells arise from the binding of cytosolic proteins to membranes. Quantifying this membrane binding and its associated diffusion in the living cell is therefore of primary importance. Dynamic photonic microscopies, e.g. single/multiple particle tracking, fluorescence recovery after photobleaching and fluorescence correlation spectroscopy (FCS) enable noninvasive measurement of molecular mobility in living cells and their plasma membranes. However, FCS with a single beam waist is of limited applicability with complex, non Brownian, motions. Recently, the development of FCS diffusion laws methods has given access to the characterization of these complex motions, although none of them is applicable to the membrane binding case at the moment. In this study, we combined computer simulations and FCS experiments to propose an FCS diffusion law for membrane binding. First, we generated computer simulations of spot-variation FCS (svFCS) measurements for a membrane binding process combined to 2D and 3D diffusion at the membrane and in the bulk/cytosol, respectively. Then, using these simulations as a learning set, we derived an empirical diffusion law with three free parameters: the apparent binding constant *K*_*D*_^app^, the diffusion coefficient on the membrane *D*_2D_ and the diffusion coefficient in the bulk/cytosol, *D*_3D_. Finally, we monitored, using svFCS, the dynamics of retroviral Gag proteins and associated mutants during their binding to supported lipid bilayers of different lipid composition or at plasma membranes of living cells and we quantified *K*_*D*_^app^ and *D*_2D_ in these conditions using our empirical diffusion law. Based on these experiments and numerical simulations, we conclude that this new approach enables correct estimation of membrane partitioning and membrane diffusion properties (*K*_*D*_^app^ and *D*_2D_) for peripheral membrane molecules.

## Introduction

The binding of cytosolic proteins to a membrane is often the starting point of dynamic processes occurring in the cell. This binding is frequently a key event in signal transduction, metabolism, membrane trafficking (endo/exocytosis) or enveloped virus assembly. Although much has been studied about these processes, quantifying the dynamics of the initial event, i.e. protein binding to and diffusion on the membrane, still remains a challenge in living cells.

Several methods, such as fluorescence recovery after photobleaching (FRAP), fluorescence correlation spectroscopy (FCS) and single(multiple) particle tracking (SPT), have been developed to monitor molecule motions in cells using fluorescence microscopy and have been extensively used in membrane biology (for review see (1)). Each of these dynamic microscopic techniques have pros and cons related to their timescale and statistics. FCS is sensitive on the millisecond to second timescale, corresponding to the diffusion characteristic time of a fluorescently-labeled molecule in a lipid mixture, through an illumination focus with a waist of approximately 200nm. FCS has therefore provided a convenient way to investigate the motions of lipids and membrane proteins in living cells. FCS usually consists in computing the auto-correlogram function (ACF) from the temporal recording of fluorescence intensity fluctuations in the illumination beam. The ACF is then fitted with an analytical expression accounting for a given type of motion e.g., fractional or normal Brownian motion in 2D or 3D. This fit yields the mobility parameters as estimates of its free parameters. Interestingly, FCS was originally developed to determine the kinetic constants of chemical reactions at equilibrium (2). Recently, FCS has been proposed as an efficient method to monitor more complex dynamics such as reaction-diffusion processes occurring in the case of transcription factor binding to DNA in cells (3) or in the embryo (4). However, single-spot FCS, in which the beam waist is constant, is usually not applicable to discriminate and quantify complex motions. In the case of reaction-diffusion processes for example, there is no analytical solution available to retrieve the kinetic parameters from the fit of the autocorrelogram. Approximated expressions can be derived by simplifying the kinetics (e.g., neglecting diffusion of the bound molecule) or distinguishing between simplification regimes (reaction kinetic dominant, diffusion dominant…) (3). However even in the case of molecules diffusing in the membrane, single spot FCS measurements often results in inaccurate estimates of diffusion coefficient, mainly because the single spot size is not sufficient to probe the heterogeneity of the environment where the molecules diffuse.

One way to circumvent that issue is to perform FCS measurements at different beam waists. Plotting the half-time of the decorrelation (*τ*_1*/*2_) as a function of the surface probed by the illumination (*w*_2_, where *w* is the beam waist, or spot size), one obtains so-called svFCS diffusion laws. Wawrezinieck et al. (5) showed that svFCS diffusion laws are a powerful tool for analyzing complex diffusions. They developed a svFCS experiment and, based on numerical simulations, managed to successfully associate svFCS diffusion laws with different types of complex, heterogeneous environments. The same approach has also been used experimentally to characterize molecular motions in membranes, below the diffraction limit (6–8) and recently in Imaging-FCS (Im-FCS) (9, 10).

In terms of FCS diffusion laws *τ*_1*/*2_ = *f* (*w*^2^), a pure Brownian motion (free diffusion) leads to *τ*_1*/*2_ *→* 0 at *w*^2^ *→* 0, i.e. the extrapolated value of *τ*_1*/*2_ for vanishing *w* is expected to be zero. Moreover, one predicts a linear relationship between *τ*_1*/*2_ and *w*^2^ with a slope that is inversely proportional to the free diffusion coefficient. Interestingly, the extrapolation of *τ*_1*/*2_ for *w*^2^ *→* 0, *τ*_1*/*2_(0) gives information about the nature of the heterogeneity probed by the molecule.

For example, molecules experiencing dynamic partitioning between liquid ordered and liquid disordered (Lo/Ld) lipid phases will have a positive *τ*_1*/*2_(0) value, whereas molecules restricted in their diffusion by sub-membrane fences, such as cortical actin cytoskeleton for example, are expected to show a *τ*_1*/*2_(0) < 0 (5, 11). However, lipids exhibiting dynamic partition in solid/liquid disordered phase (S/Ld) also display *τ*_1*/*2_(0) < 0 (12), showing that the *τ*_1*/*2_(0) value alone is not sufficient to correctly characterize the motion (13).

Recently, different types of svFCS diffusion laws have been derived and characterized for different types of heterogeneous environments based on numerical simulations (14, 15), mostly for complex motions related to anomalous subdiffusion. In this article, we explored the applicability of svFCS diffusion laws to the quantification of membrane binding kinetics where the fluorescent molecule diffuses both at the 2D membrane and in the 3D bulk. Our main objective was to test the suitability of such experimental diffusion laws to quantify the main parameters: the apparent binding constant (*K*_*D*_^app^), the membrane bound diffusion coefficient (*D*_2D_) and the cytosolic/bulk diffusion coefficient (*D*_3D_) (see table 1 for the definitions of the different parameter use here).

**Table 1.** List of parameters used in this work

| Variables | Parameters details |
| --- | --- |
| *Confocal volume calibration parameters* |  |
| $w_{xy}$ | Width of the observation area in x, y direction |
| $s = w_z/w_{xy}$ | Shape factor of the 3D Gaussian excitation point spread function (PSF) |
| *Parameters used in ACF fit* |  |
| $G(\tau)$ | Autocorrelation function (ACF) |
| $\tau_{1/2}$ | Characteristic time of half temporal decorrelation i.e.<br>$G(\tau_{1/2}) = (G(0) + G(\infty))/2$ |
| $\tau_d$ | Characteristic diffusion time of particles in the confocal volume<br>$\tau_d = \tau_{1/2}$ for 2D diffusion |
| $N$ | Effective number of particles in confocal volume |
| $T$ | Fraction of particles in triplet state |
| $\tau_T$ | Characteristic residence time in triplet state |
| *Parameters of the 2D/3D diffusion and binding kinetics* |  |
| $D_{2D}$ | Diffusion coefficient of membrane-bound molecules |
| $D_{3D}$ | Diffusion coefficient of free molecules |
| $k_{on}$ | Association constant of particles to the membrane |
| $k_{on}^* = k_{on}c$ | Apparent association constant of particles to the membrane<br>(depends on the binding target concentration, $c$) |
| $k_{off}$ | Dissociation constant of particles from the membrane |
| $K_D = k_{off}/k_{on}$ | Binding affinity of the particles to the membrane |
| $K_D^{app} = k_{off}/k_{on}^*$ | Experimental apparent binding affinity of the particles<br>to the membrane (depends on the binding target concentration, $c$) |

We first performed extensive numerical simulations of svFCS diffusion laws for this system, exploring different regimes by varying *K*_*D*_, *D*_2D_ and *D*_3D_. Using part of these simulations as a learning set, we could derive an empirical svFCS diffusion law and use the other part of the simulations as a test set to confirm that the diffusion law can indeed be used to estimate *K*_*D*_ and *D*_2D_ of synthetic data with good accuracy. We then used this empirical diffusion law to fit experimental svFCS measurements of HIV-1 or HTLV-1 retroviral Gag proteins binding to either supported lipid membranes or living HEK 293T cells plasma membranes. HIV-1 and HTLV-1 Gag proteins are multi-domain proteins binding the plasma membrane of host cells and are essential for viral assembly. HIV-1 Gag for example contains three main domains, each having distinct roles during viral assembly. The matrix domain (MA) is responsible for membrane binding to the PI(4,5)P_2_ lipids, present in the inner leaflet of the plasma membrane, thanks to a highly basic region (HBR). Membrane binding is reinforced by the N-terminal myristate MA substitution that inserts into the inner leaflet. The capsid domain (CA) is involved in Gag-Gag interactions during Gag self-assembly, the initial stage in the generation of a new virion. The nucleocapsid domain (NC) also contains enriched basic motifs that are involved in RNA binding to permit encapsidation of the RNA in the new virion. NMR data and coarsed grain molecular dynamics have shown that the HBR region of MA as well as other polybasic motifs bind to anionic lipids (PI(4,5)P_2_, but also PS) (16–19). Conversely, the lack of myristate has been shown to decrease membrane binding of HIV-1 in living cells (20). Finally, Fogarty et al. (21) have shown that HTLV-1 Gag has a higher affinity for cell plasma membranes than HIV-1 Gag.

Although these results from the literature do not consist in a precise quantification of the binding affinities, they provide us with ordering relations between them, e.g. the binding affinity of HIV-1 in the presence of myristate is higher than in its absence. We used them to challenge the capacity of our svFCS diffusion law to retrieve the correct estimations from experimental svFCS diffusion laws and to correctly estimate *K*_*D*_^app^ and *D*_2D_ *in vitro*, using supported lipid bilayers (SLBs) of various composition and *in cellulo* by expressing Gag chimerae proteins in HEK-293T cells.

On the synthetic test dataset, we show that our svFCS diffusion law yields precise estimates for *D*_2D_ and *K*_*D*_, with an accuracy of 14% and 24%, respectively. The accuracy of our method on real experimental measurements cannot be assessed directly, but our results confirm the capacity of our 2D/3D diffusion and binding svFCS diffusion law to correctly estimate *D*_2D_ and *K*_*D*_^app^ in the case of HIV-1 and HTLV-1 Gag proteins binding to model lipid SLB and to HEK-293T cells. Notably, our estimates confirm previous results obtained in the literature with other methods by different teams. Therefore, our results highlight the benefits of our empirical svFCS diffusion law for the determination of *D*_2D_ and *K*_*D*_^app^ in the membrane binding problem with diffusion both at the membrane and in the cytosol.

## Materials and Methods

### Lipids and Plasmids

Egg-PhosphatidylCholine (PC), Brain-PhosphatidylSerine (PS), Brain-Phosphatidyl Inositol (4,5)BisPhosphate_2_ (PI(4,5)P_2_), 1,2-dioleoyl-sn-glycero-3-phosphoethanolamine-N-(Cyanine 5.5) (PE-Cy5.5) and 1-palmitoyl-2-(dipyrrometheneboron difluoride)-undecanoyl-sn-glycero-3-phosphocholine (TopFluor®-PC) were purchased from Avanti Polar Lipids (Alabama,USA). Atto647N-PI(4,5)P_2_ is a gift from Pr. Christian Eggeling (Jena, Germany). The plasmid encoding HIV-1-myr(-)Gag-GFP is a gift of Dr. H.de Rocquigny and described in (22). The plasmid encoding HTLV-1 Gag-YFP (named pHTLV-Gag-YFP) was obtained from Dr. D. Derse’s lab and described previously in (23). The pHIV-1-Gag-mCherry was obtained from Dr. N. Jouvenet (24).

### Gag purification and labelling

Purified Gag Proteins were produced and provided by Pr. J. Mak’s lab (25). Protein stock concentration was measured at 1.2mg/mL using a NanoPhotometer (Implen), and 100*µ*L were incubated overnight under agitation and at 4°C with 1*µ*L of Alexa Fluor 488 C5-Maleimide (Invitrogen) at 20mM in DMSO. The reaction mix was transferred in Slide-A-Lyzer MINI Dialysis Device, 0.5mL (Thermo Scientific) and dialyzed 6h under agitation at 4°C in 15mL Buffer; TRIS (50mM), NaCl (1M), pH=8. Dialysing buffer was then renewed, and the reaction mix was dialysed again overnight under agitation at 4°C, collected, and stored at -20°C.

### LUVs processing

Liposomes were processed by dissolving in chloroform the following lipids aiming at a 1mg/mL concentration: Egg-PC, Brain-PS, Brain-PI(4,5)P_2_ and PE-Cy5.5 in the ratios described Figure 2a. Solvent was then evaporated during 20min in a rotary evaporator and 10min more in a desiccator. Lipids were rehydrated in 500*µ*L Na Citrate Buffer; Na Citrate (10mM), NaCl (100mM), EGTA (0.5mM), pH 4.6. The mixture was then, 5 consecutive times: frozen 30s in liquid nitrogen, water bathed at 37°C for 30s and vortexed. MLVs processed as such can be stored at - 20°C. LUVs are subsequently produced by diluting Liposomes 1:5 in Na Citrate Buffer. This solution is then passed through a 100nm Nucleopore Track-Etched Membranes in an extruder (Avanti Polar Lipids), and sonicated.

### LUV binding experiments

Proteins at a final 50nM concentration were incubated 30min at room temperature with LUVs compositions described Figure 2a at a final 0.8mg/mL concentration. 100*µ*L samples were then centrifuged at 42000rpm for 30min at 4°C (Beckman TLA 100 rotor). 2 fractions were isolated from these samples : 90*µ*L were collected from the supernatants, and the pellets were resuspended with 80*µ*L TRIS NaCl Buffer; TRIS HCl (10mM), NaCl (150mM), pH 7.4. Equal volumes from each fraction were loaded in SDS-PAGE gels and analysed by Western Blot. Gag proteins were detected using an anti-p24 as primary antibody and a secondary antibody coupled to HRP (Horse Radish Peroxydase). Membranes were imaged by enhanced chemiluminescence (ECL) in the ChemiDoc (Biorad Inc, USA) and bands relative intensities were quantified using ImageJ. *K*_*D*_^app^ values were then calculated as in equation 1, ie the ratio of the bands signal corresponding to supernatants (*S*) over the bands signal corresponding to pellets (*P*) corrected by the pellet obtained with Gag alone *P*_*GA*_, which account for spontaneous self-assembly. According to the mass action law, the amount of self-assembly was considered proportional to the total amount of Gag (*T* = *P* + *S*) and normalize to this total amount when Gag was alone (*T*_*GA*_). To correctly estimate *K*_*D*_^app^ we then performed the following calculation :

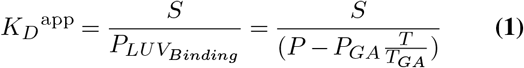

### SLBs processing

Glass coverslips (25 mm No.1.5 H cover glass (Marienfeld)) were treated 30min with ozone in Ossila UV Ozone Cleaner and rinsed thoroughly with ultrapure water. The sample is delimited by a 7mm diameter plastic cylinder fixed to the coverslip using Twinsil (Picodent). 100*µ*L of 0.2mg/mL SUVs of the different mix compositions are spread on the coverslip and incubated 40min at 37°C. Formed SLBs are washed 4 times with filtered TRIS NaCl Buffer to remove non-fused vesicles.

### Living cell sv-FCS experiments

Human embryonic kidney cell line (HEK-293T) were seeded on 25 mm No.1.5 H cover glass (Marienfeld) coated with poly-L-lysine (Sigma). 0.15 million cells were maintained in 2ml Dulbecco’s Modified Eagle’s Medium (DMEM, GIBCO) for 24 hours at 37°C. Transfection was performed using CaCl_2_ (250mM) and HBS2X, with 1 *µ*g pHIV-Gag-mCherry, 0.3 *µ*g HIV-GagG2A-GFP and 1*µ*g pHIV-Gag-YFP. 15 hours post transfection, medium was replaced with phenol-red free medium L15 (GIBCO) supplemented with 20 mM Hepes, for sv-FCS experiments on live cells.

### Acquisition and fit of the experimental diffusion laws

*FCS set up* FCS was performed on a Zeiss LSM 780 microscope equipped with the variable pupil coverage system using water immersion (NA = 1.2). The underfilling of the objective back - aperture leads to the increasing of the beam waist. *Beam waist calibration w*_2_ was calibrated using highly diluted Rhodamine-6G (R6G) and Tetramethylrhodamine (TMR, a gift from Dr. M.May) solutions excited at 488 nm and 561 nm respectively. At least 30 autocorrelogram functions (ACF) were obtained at T=37*°*C from 5S fluctuation intensity acquisition. Laser irradiance of the sample is adjusted for the different waists in order to get constant molecular brightness. The diffusion coefficient of R6G and TMR solutions are set to D = 298 *µ*m^2^.s^*−*^1 and the free diffusion time is obtained by fitting the autocorrelation function of the measured time - trace intensity with an analytical model of a 3D free diffusion including a triplet model (T + 3D).

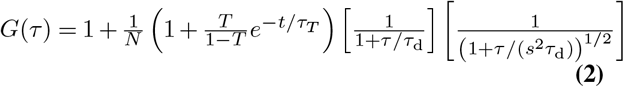

Rhodamine triplet time was measured from the smallest waist correlogram to be 4*µs*, and was therefore kept fixed to fit all the bigger waist calibration correlogram, only T, the triplet fraction was allowed to change. *s* is set to 5. We measured excitation waists varying from 170 to 700 nm (488nm excitation) or from 210 to 800 nm (561 nm excitation) depending on objective’s back-aperture coverage. *FCS diffusion law acquisition on SLBs and cell* The SLB z=0 axial position was retrieved thanks to the Cy5.5 PE fluorescence, ACF were recorded at 25°C. Cell intensity fluctuations and associated ACF correlograms were acquired at 37°C by parking the laser in the z-axis at the level of the bottom membrane, in order to avoid deformation of the laser shape due to index mismatch that will occur at the top membrane. The position of the laser spot into the cell was systematically controlled with a z-scan intensity profile before and after each waist of correlogram acquisition. In both cases, FCS measurements were done for 7 different beam waist and we recorded at least 50 series of time-trace intensity of 10s duration per waist by successive series of 10. Again,the laser irradiance is adjusted over the different waists to keep constant and independent of the waist probed a significant photon count per molecules in the range of 2-5 kHz, but also kept low enough to avoid important photobleaching. This is controlled *a posteriori* by plotting *N*, the number of molecules as a function of *V*_*eff*_ = *π*^3/2^*sw*^3^ (see Fig.SI.2). In the case of GFP and YFP labelled proteins as well as for the Alexa-Fluor 488 labelled Gag, each ACF correlogram function are fitted (from 10*µ*s to 1.7s) with an analytical model for Brownian 2D diffusion.

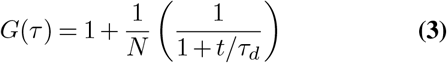

FCS measurements of mCherry labelled proteins were done with a 561 nm laser line. mCherry is known to exhibit flickering which lead to dark state population with typical ground state recovery kinetics (26). As we didn’t established an analytical model for this particular photophysics, we arbitrarily took this phenomenon into account in our fit by splitting our 2D diffusion ACF expression into two components with equivalent apparent molecular brightness :

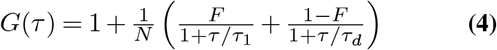

with *τ*_1_ << *τ*_*d*_. As we always use similar laser irradiance amongst the different experiments, we observed 0.45 < *F* < 0.65 when fitting the ACF data between 0.2*µ*s and 1.7s.

## Numerical simulations of (sv)FCS experiments

### Algorithm

We emulated FCS experiments by simulating the binding reaction and diffusion of 100 individual molecules in a cubic nucleus of size *L*^3^ *µ*m^3^ during 11 seconds. Unless otherwise indicated, we use *L* = 6*µ*m in this work. The plasma membrane was defined as the *z* = 0 face of the cell, with the cytoplasmic bulk as the *z* > 0 3D volume. At the start of the simulation, a fraction of the individual molecules were positioned at random 3D positions (uniform distribution) within the cytoplasm and the reminder was located at random 2D positions (uniform distribution) on the plasma membrane (i.e. with coordinates *z* = 0) (Fig 1a1). The fraction of molecules initially free in the cytoplasm was set to its theoretical equilibrium fraction i.e., to *k*_off_ */*(*k*_on_ + *k*_off_). Reaction-diffusion simulation processed by successive iterations of time step Δ*t* (we used Δ*t* = 1*µ*s). The position **r**_*i*_ of every individual molecule *i* = {1,…, 100} was first moved according to independent Brownian dynamics: 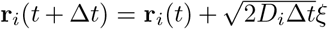 where *ξ* is a 3D vector of i.i.d. Normal random numbers and *D*_*i*_ = *D*_2D_ or *D*_3D_, for membrane-bound or free molecules in the bulk, respectively. Membrane-bound molecules were kept at the membrane at this step (i.e. we reset their coordinate *z* to zero).

**Fig. 1.**
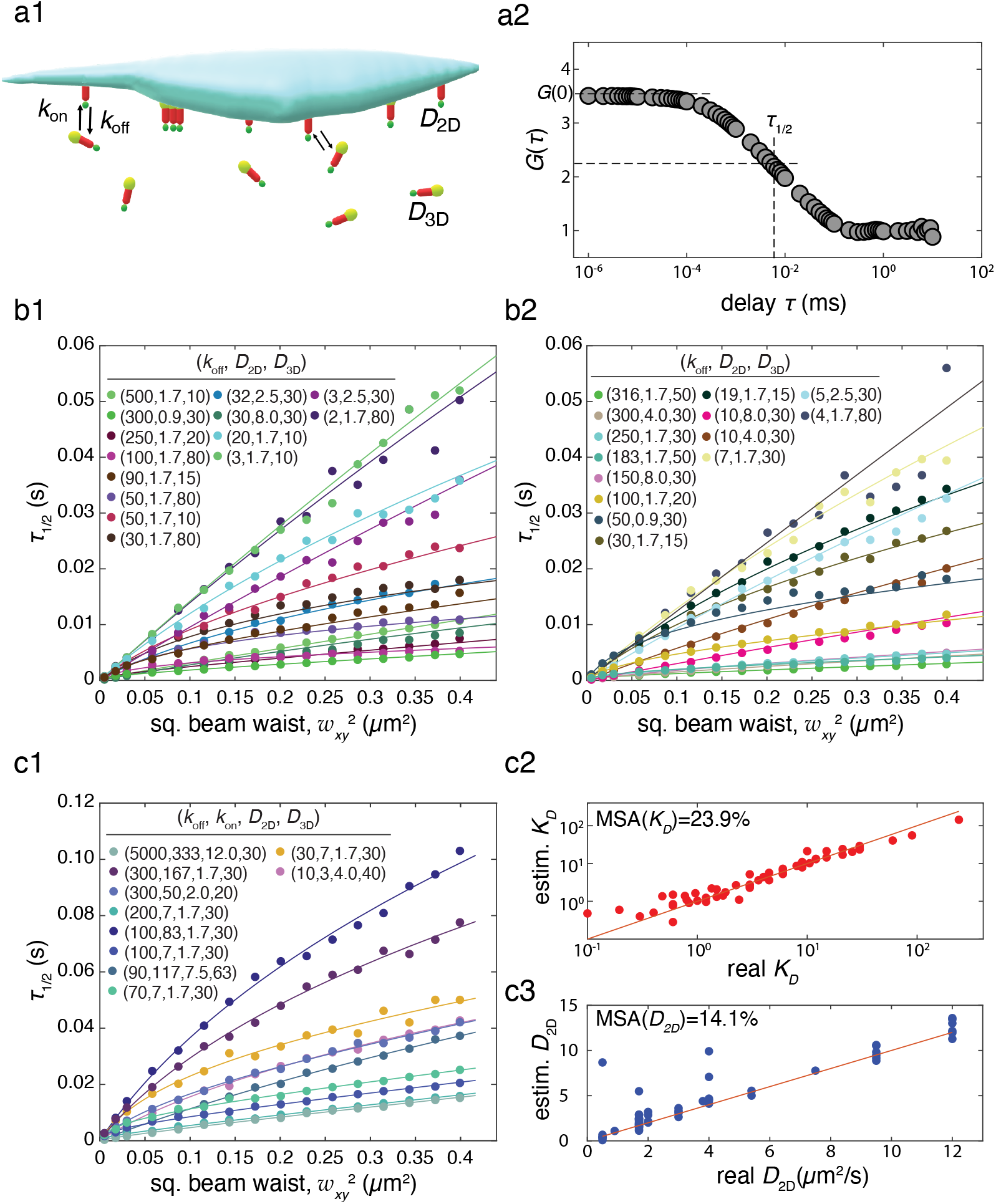
Simulation-based derivation of a phenomenological diffusion law : a1- We used the numerical simulations of svFCS described under Material and Methods to generate synthetic data of an svFCS experiment where HIV-1 gag molecules diffuse freely in 3D in the bulk / cytoplasm or in 2D as bound molecules at the plasma membrane, with binding and unbinding events driven by rate constants *k*_on_ and *k*_off_, respectively. a2- The algorithm outputs the corresponding auto-correlation function, that we quantify by its FWHM (full width at half maximum), *τ*_1*/*2_. b- The diffusion laws, i.e. the changes of *τ*_1*/*2_ with beam width *w*_*xy*_^2^, are first fitted with the phenomenological diffusion law eq.(6) to the “training set” in order to estimate the six fit parameters of the law (see text). We show a selection of the synthetic diffusion laws (full circles) and their fits (full lines). The values of *k*_off_, *D*_2D_, *D*_3D_ for each simulation are given in the legend. For the training set, we fix *k*_on_ = 16.67s^*−*1^. c1- Fitting eq.(6) to the synthetic data of the test set is then used to estimate *k*_on_, *k*_off_ and *D*_2D_ (*D*_3D_ is considered known, see text). The panel illustrates the obtained fits for a randomly chosen sample of simulations. c2 & c3- The estimated values of *K*_*D*_ = *k*_off_ */k*_on_ or *D*_2D_ are compared to their real values. The accuracy of these estimations is quantified using the Median Symmetric Accuracy (MSA).

Classical boundary conditions e.g., periodic or reflective, introduce strong correlations in the simulations that are not present in the experiments. We therefore opted for the following scheme. Molecules exiting the cell through the plasma membrane at *z* = 0 were changed into bound molecules (thus reset at *z* = 0), whereas the upper face (*z* = *L*) was considered reflective. Molecules exiting the cell through any of the 4 remaining faces were removed from the simulation. To keep the number of molecules constant in the cell, removed molecules were compensated by the addition of an equal number of new free molecules that were located in the bulk at a distance *ϵ* = 10 nm of one of the 4 remaining faces, chosen at random (with uniform distribution).

We then simulated the reaction step. We considered that free bulk molecules located at a distance less than *ϵ* = 10 nm from the plasma membrane (i.e., for which *z* < 0.01*µ*m) were close enough to bind. Each of these molecules was therefore independently turned into a membrane-bound molecule with probability *p*_on_. Conversely, every membrane-bound molecule was independently allowed to unbind i.e., was changed into a free bulk molecule, with probability *p*_off_. Both reaction probabilities were set from the simulation parameters according to: *p*_on_ = *k*_on_*L*Δ*t/ϵ* and *p*_off_ = *k*_off_ Δ*t*. To emulate fluorescence emission, we used a 3D Gaussian illumination profile centered on the plasma membrane at *z* = 0. The probability that a molecule located at position (*x, y, z*) at time *t* emit one photon between *t* and *t* +Δ*t* was computed as 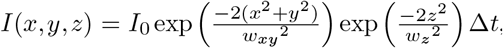, with *w*_*xy*_ and *w*_*z*_ the beam width in the *x, y* or *z* direction respectively. We set *I*_0_ to 15 *×* 10^4^ to reproduce the experimentally-observed brightness. To achieve correct equilibrium in the molecule locations before recording photon emission, the collection of photon emission times was switched off during the first 500 ms of every simulation run. We counted the total number of photons emitted at each Δ*t* time step during the simulation before computing the auto-correlation of the corresponding time series. For each parameter value, we ran 20 independent realisations of this 11 second-simulation, and averaged the 20 resulting auto-correlations to get the average auto-correlation *G*(*τ*).

### Simulation of spot variation FCS

To emulate an svFCS experiment, we repeated the above simulations for a selection of 16 values of the beam width *w*_*xy*_ between 71 and 623 nm and obtained the auto-correlation *G*(*τ*) for each of these 16 values. The beam width in the *z* direction was set to *w*_*z*_ = *sw*_*xy*_ with a constant value *s* = 4.3 (a crude estimate from the experimental setup yields *s ≈* 5).

### Code availability

The computer code used in this article is freely available at https://gitlab.inria.fr/hberry/gag_svfcs.

## Results

### Empirical determination of a phenomenological diffusion law

#### Single spot size

A first possibility to interpret the results of FCS experiments is to derive an analytical expression for the auto-correlation function *G*(*τ*) and fit it to the data to estimate the parameters. In practice, this is possible only for the simplest cases. In particular, this is not feasible, to our knowledge, when the problem is not homogeneous nor isotropic as is our case here with a 2D membrane located in a 3D volume at *z* = 0. However, one can simplify the problem by considering that the membrane binding and diffusion sites are homogeneously distributed in the 3D bulk, and by neglecting the reduction of dimensionality. The result is a system where the bound molecules diffuse in the same 3D space as the free ones i.e., a spatially homogeneous reaction-diffusion problem with reaction 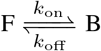 with *F* the free molecules diffusing with diffusion coefficient *D*_3D_ and *B* the bound molecules diffusing with diffusion coefficient *D*_2D_. Textbook reviews and several articles have treated simplified versions of the problem. For instance, the correlation function for the case *D*_2D_ *≈* 0 is derived in Michelman-Ribeiro et al. (3) whereas Krichevsky and Bonnet (27) derives it for *D*_3D_ = *D*_2D_. However, we could not solve the general case *D*_3D_ > *D*_2D_ > 0 (see Supporting Information SI1).

An alternative approach consists in fitting the autocorrelation curves *G*(*τ*) = *f* (*τ*) with a selection of expressions derived for other problems and test on synthetic (simulation) data the accuracy or meaning of the parameters thus estimated. Supporting Information Figure SI.1 shows the result of this approach. Here, we simulated diffusion and binding for a range of *k*_on_ and *k*_off_ values using the algorithm described above with *D*_2D_ = 1.7 and *D*_3D_ = 30 *µ*m^2^*/*s. For each parameter value, we fitted the obtained auto-correlation function with three models: a single population with 3D Brownian motion,

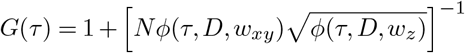

with *φ*(*τ, D, w*) = 1 + 4*Dτ/w*^2^, a mix of two populations with 3D Brownian motion,

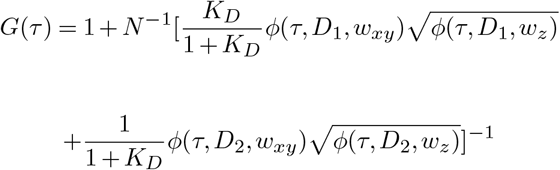

or anomalous diffusion

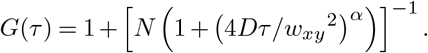

The best-fit model was then selected using corrected Akaike information criterias (28) (Fig SI.1a). We found that the best-fit model is the single Brownian model with low diffusion coefficient for *k*_on_ *≫ k*_off_, or with large diffusion coefficient for *k*_off_ *≫ k*_on_. Therefore, in these regimes, the approach correctly spots one of the two diffusion coefficients, but is blind to the other. Between these extreme regimes, the best-fit model was found to be the 2 population model with one small and one large diffusion coefficients (Fig S I.1b1). For most of the (*k*_on_, *k*_off_) values, this simple procedure is remarkably precise regarding the estimation of the diffusion coefficients (Fig SI.1b2). However, the estimation for *K*_*D*_ is very bad, with estimated values that can be one order of magnitude smaller than the real value (Fig SI.1b3). This was of course to be expected, given the simplifying hypotheses that support this approach. However, since our experimental interest here is mainly on the estimation of *K*_*D*_, we opted for a strategy based on spot variation.

#### Spot variation FCS

Our motivation for svFCS is based on the following argument. We consider here a FCS setup where the PSF is centered on *z* = 0 at the plasma membrane. Let *F* and *B* be the concentration (or density) of molecules in the cytoplasm or bound at the membrane, respectively. Then the number of bound molecules found inside the focal volume is *N*_*B*_ *∝ Bw*_*xy*_^2^, whereas the number of free molecules found in the focal volume is *N*_*F*_ *∝ Fw*_*xy*_^2^*w*_*z*_ = *F sw*_*xy*_^3^. The ratio between both is thus *N*_*B*_*/N*_*F*_ = *B/F ×* 1*/*(*sw*_*xy*_). The concentrations *B* and *F* are constant. The ratio *s* = *w*_*z*_*/w*_*xy*_ is as well roughly constant in our experimental setup. Therefore we expect from this simple analysis that the svFCS signal at very small beam widths *w*_*xy*_ will mostly be dominated by the bound molecules whereas free molecules should dominate at very large spot sizes.

In the svFCS literature, the evolution of *τ*_1*/*2_,the FWHM of *G*(*τ*) (full width at half maximum i.e., the value of *τ* were *G*(*τ*) is half its maximum value, see Fig 1a2) is referred to as a diffusion law. A first phenomenological diffusion law for our case therefore consists in expressing *τ*_1*/*2_ as the sum of a 2D and a 3D contribution, under the constraint that the 2D bound fraction should dominate, i.e. *τ*_1*/*2_ *→ w*_*xy*_^2^*/*(4*D*_2D_) for *B/F→ ∞* or *w*_*xy*_ *→* 0, whereas the 3D free fraction should dominate i.e., *τ*_1*/*2_ *→ w*_*xy*_^2^*/*(4*D*_3D_) for *B/F* → 0 and *w*_*xy*_ *→ ∞*. A simple ansatz that respects these constraints is:

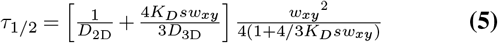

Unfortunately, eq.(5) was not found to yield correct estimates of *D*_2D_ or *K*_*D*_ on synthetic data. We therefore generalized it to a more complex phenomenological expression:

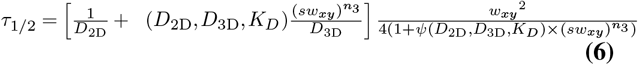

with

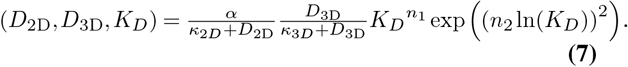

In addition to the simulation parameters (*D*_2D_, *D*_3D_, *K*_*D*_, *s*), eq.(6) comprises six free fit parameters: *α, n*_1_, *n*_2_, *n*_3_, *κ*_2*D*_ and *κ*_3*D*_. We next estimated their values using a simulation-based approach.

We ran a total of 160 svFCS simulations, each with different values for *D*_2D_, *D*_3D_, *k*_on_ and *k*_off_, thus obtaining 160 diffusion laws *τ*_1*/*2_ = *f* (*w*_*xy*_^2^). These 160 simulations were distributed into two groups: for 114 simulations (“training set”), we kept the same value for *k*_off_ (16.67 s^*−*1^) while the three remaining parameters were varied; for the 56 remaining simulations (“test set”), all the four parameters were varied. No parameter values were common between the two sets.

In a first stage (“training”), we considered the values of *D*_2D_, *D*_3D_, *k*_on_ and *k*_off_ as known, and fitted the 114 diffusion laws of the training set with *α, n*_1_, *n*_2_, *n*_3_, *κ*_2*D*_ and *κ*_3*D*_ as free parameters. Figure 1b gives a random selection of the diffusion laws and the fits that were obtained. The resulting values for the parameters of the phenomenological diffusion law eq.(6) were: *α* = 0.668, *κ*_2*D*_ = 0.270, *κ*_3*D*_ = 13.184, *n*_1_ = 1.115, *n*_2_ = 0.236 and *n*_3_ = 1.532.

In a second stage, we fixed the values of *α, n*_1_, *n*_2_, *n*_3_, *κ*_2*D*_ and *κ*_3*D*_ to their above values and fitted the phenomenological diffusion law eq.(6) to the 56 diffusion laws of the test set using *D*_2D_, *k*_on_ and *k*_off_ as free fit parameters. We considered that *D*_3D_ can easily be determined from independent experiments (see below) and therefore considered it as known. Note that the values of *D*_2D_, *D*_3D_, *k*_on_ and *k*_off_ used for this test set have never been used in the training set. A random choice of the obtained fits is shown in Figure 1c1. For each of these 56 simulations, we compare the estimates of *K*_*D*_ = *k*_off_ */k*_on_ and *D*_2D_ thus obtained with their real values on Fig. 1c2 and c3, respectively. In these panels, the main diagonals (red line) represent perfect estimations. In both cases, the estimation is most of the time close to the diagonal, indicating a good estimation. Although *K*_*D*_ was varied over more than 3 orders of magnitude, its estimation is correct, at least in the range [0.3,30]. Concerning *D*_2D_, except for a handful of over-estimations, the quality of the estimation is also rather acceptable. We quantified the accuracy of these estimates by the median symmetric accuracy (29): MSA = 100exp (*M* (|ln(*y*^pred^*/y*^real^)|) −1), where *M* designates the median function and *y*^pred^ or *y*^real^ are the predicted or real values of the parameters. We found an estimation error of 24 % and 14 % for *K*_*D*_ and *D*_2D_, respectively. With this accuracy, the phenomenological diffusion law eq.(6) can clearly not be considered a very precise method to estimate *K*_*D*_ and *D*_2D_. However, we considered it provides us with estimates that are correct enough to be tested using experimental data.

### Monitoring HIV-1 Gag partition using FCS diffusion laws on model membranes

To assess the ability of our empirical analytical expression of FCS diffusion law (eq.6) to correctly estimate the *K*_*D*_^app^ and the membrane bound diffusion coefficient *D*_2D_ in the case of a 2D/3D binding unbinding kinetics, we performed svFCS of HIV-1-myr(-)-Gag protein kinetics in the presence of supported lipid bilayers (SLBs) as depicted in figure 2a. HIV-1-Gag is known to interact with lipid membranes thanks to a bipartite motif consisting of a myristate and a polybasic domain (Highly basic region, HBR) (for review see (30)). Electrostatic interactions of HIV1-Gag with negatively charged lipids (such as phosphatidylserine (PS) and phosphatidylinositolbisphosphate (PIP_2_) have been shown using NMR (17, 18).

**Fig. 2.**
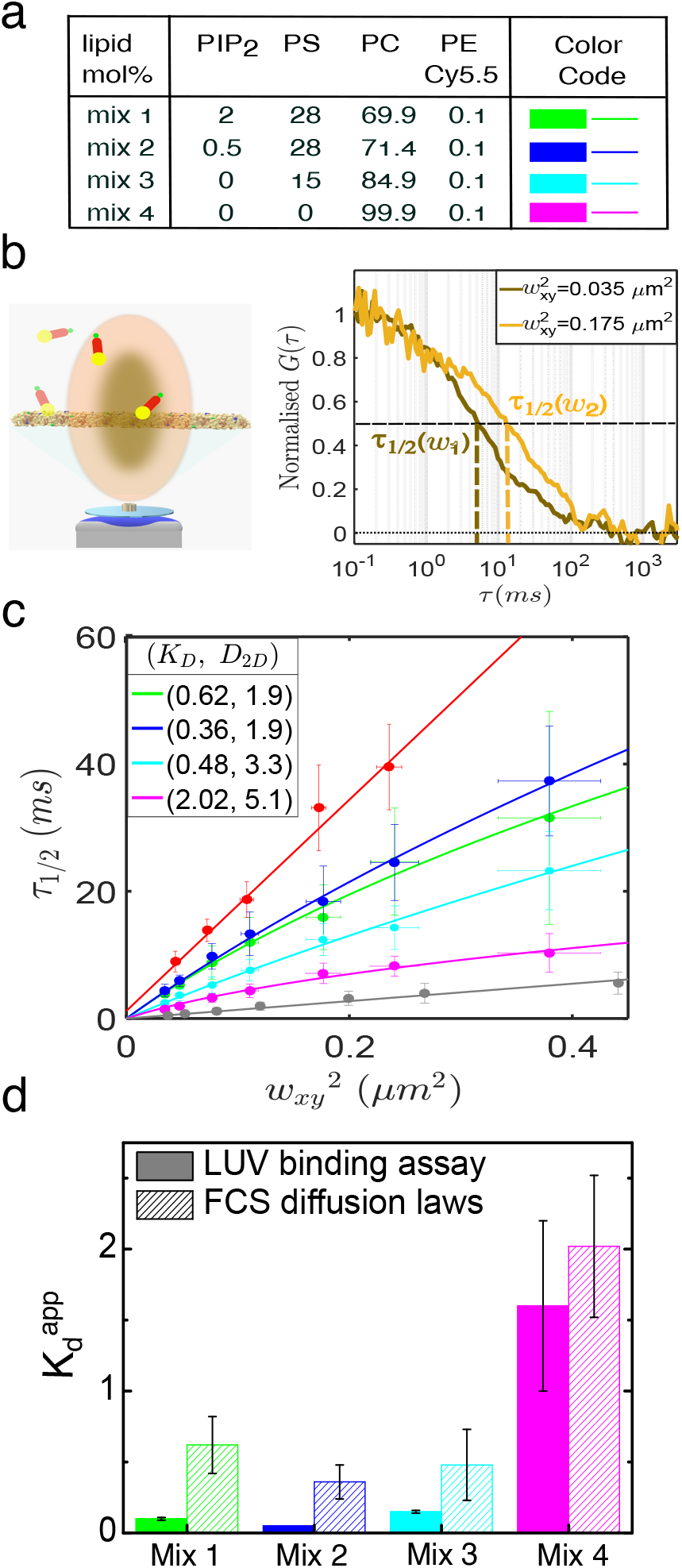
FCS diffusion laws on model membranes : a- Different composition of the SLB used here. b- These SLBs were spread on glass coverslips and set in the microscope to perform sv-FCS measurements. Fluorescent myr(-)Gag was introduced in the buffer at the beginning of the experiment. Example of normalized correlograms recorded during an experiment at two different laser waists (190 nm and 420 nm) showing increasing *τ*_1*/*2_ values with increasing waists. In order to establish the diffusion laws, the *τ*_1*/*2_ values are obtained at different waists by fitting the correlogram with equation 3. c- Experimental FCS diffusion laws of myr(-)Gag obtained at the surface of the different SLB composition (dots) and their fits (lines) using eq.6 with *K*_*D*_^app^ and *D*_2D_ as free parameters (see table for colour code). In grey, FCS diffusion law of myr(-)Gag in the buffer (dots) and its fit using a linear model (line). In red, FCS diffusion law of Atto647N-PIP_2_ in the supported lipid bilayer (dots) and its fit with a linear model (line). d- Comparison of the *K*_*D*_^app^ values obtained from LUV binding assay (full bars) and sv-FCS experiments fits (dashed bars). Values are mean*±*sd obtained from the fit in the case of sv-FCS and mean*±*sd of n=2 LUV binding experiments.

We monitored differences occurring in membrane binding using four different lipid compositions containing decreasing amount of negatively charged lipids (namely mix 1 to 4, see the table in figure 2a for the composition and the molar proportion of each lipid). Interaction with the plasma membrane is a key step in the initiation of HIV-1 Gag self-assembly, occurring during the generation of a new virion (31). However, HIV-1 Gag self assembly only occurs above a critical concentration of HIV-1 Gag. Yandrapalli et al. (32) shown that for HIV-1 Gag concentration below 50 nM, no self-assembly was observed on SLB with composition equivalent to our mix1. To avoid self-assembly, in order to stay in the limit of our kinetic model, fluorescent HIV-1 Gag was injected in the bulk phase, above the SLB, to a final concentration of 10 nM. In parallel, fluorescent PE-Cy5.5 was added to a negligible molar proportion into the lipid composition in order to precisely locate the SLB in the axial direction and correctly focus the laser to ensure maximal *w*_*xy*_ value at the SLB. Correlograms were then collected at different waists, as illustrated in figure 2b to determine the mean value of *τ*_1*/*2_ for each waist, in order to establish the svFCS diffusion laws. Figure 2c shows the svFCS diffusion laws (*τ*_1*/*2_ = *f* (*w*_*xy*_^2^)) obtained for the four different lipid composition, from the highest charged one (mix 1, blue dots) to the neutral one (mix4, pink dots). Each *τ*_1*/*2_ value plotted on these diffusion laws represent the mean *±* sd of 30 < *n* < 110 correlograms measured on 2 < *n* < 5 different SLBs for each *w*_*xy*_^2^.

Each *w*_*xy*_^2^ value is itself a mean *±* sd of 60 different measurements on rhodamine6G standard. Interestingly, from figure2c, it can be seen that each of these experimental diffusion laws are flanked by two different diffusion laws. One, the red dots, is the diffusion law of Atto647N-PI(4,5)P_2_ inserted in the mix1 SLB with its linear fit for free diffusion model (red line). The other, in gray dots, was obtained by measuring *τ*_1*/*2_ decorrelation times of labelled myr(-)Gag in the buffer, far away (z= 15*µ*m) from the mix1 SLB, with its linear fit for free diffusion (gray line). These linear fits lead to the following diffusion coefficients: in the case of Atto647N-PI(4,5)P_2_, *D*_2D_=1.9 *±* 0.2 *µ*m^2^.s^*−*1^ and in the case of labelled myr(-)Gag, *D*_3D_=19 *±* 2 *µ*m^2^.s^−1^. In comparison, each of the diffusion laws obtained for labelled myr(-)Gag when focusing the laser at the SLB does not seem to have a linear tendency. On the opposite, depending on the lipid composition, they exhibit different curvature as in our simulated diffusion laws (see fig.1b1,b2,c1), suggesting a 2D/3D diffusion plus binding/unbinding process. These diffusion laws were fitted with eq.(6), in order to extract *D*_2D_, *K*_*D*_^app^ and *D*_3D_ values. Fitting the experimental diffusion laws leaving the three parameters free, systematically led to highly erroneous *D*_3D_ (see figSI.2 for details). For this reason, we fixed *D*_3D_=19 *µ*m^2^.s^*−*1^ in our fit, as experimentally determined. In this case, we measured *D*_2D_ = 1.9 *±* 0.2 *µ*m^2^.s^*−*1^ for myr(-)Gag in the case of PI(4,5)P_2_ containing SLBs (mix 1& 2). Interestingly this value is similar to the diffusion coefficient value measured for Atto647N-PI(4,5)P_2_ in mix 1 using Brownian svFCS diffusion laws. These *D*_2D_ then have higher values in SLBs lacking of PI(4,5)P_2_ and increase with decreasing surface charges from 3.3 to 5.1 *µ*m^2^.s^*−*1^. These determinations therefore are in perfect agreement with the previous literature on the effect of PI(4,5)P_2_ on myr(-)Gag.

However, our main goal here was to assess the ability to measure *K*_*D*_^app^s using svFCS. To compare our quantification with results independently obtained with a standard method, we performed LUV binding experiments with the 4 different lipid mixtures of figure2 (see also supplemental figSI.2). Figure 2d shows the *K*_*D*_^app^ values obtained respectively by LUV binding experiments (full bars) and by fitting the FCS diffusion laws (dashed bars) with our empirical expression eq.(6). For every lipid mixture, the *K*_*D*_^app^ determined by our empirical diffusion eq.6 were systematically larger than the those obtained by LUV binding assays. However, overall, our estimation of *K*_*D*_^app^ with the empirical svFCS diffusion law follows exactly the same trend than thise obtained with LUV binding assay. Only mix4, that has no negatively charged lipids, exhibits a *K*_*D*_^app^ > 1, i.e. a partition of myr(-)Gag in favor of the bulk phase, whereas all the other composition (mix1 to mix3) are found to have myr(-)Gag mainly partitioning at the lipid membrane (*K*_*D*_^app^ < 1).

### Quantifying different retroviral Gag proteins binding at the plasma membrane of HEK-293T living cells

We then explored the capability of our svFCS experiments to measure *K*_*D*_^app^ and *D*_2D_ values in living cells. To this aim, we transfected HEK-293T cells with plasmids expressing either HIV-1-Gag-mCherry or HIV-1-myr(-)GagGFP or HTLV1-GagYFP chimera proteins. Figure 3a shows representative confocal fluorescence microscopy images of the HEK-293T cells expressing these three proteins. As can be seen from the fluorescence intensities in the images, fluorescent protein expression amongst the cells was heterogeneous. We selected the cells with the lowest fluorescence intensity to perform the sv-FCS experiments as HTLV-1 (poorly) as well as HIV-1 Gag (mostly) proteins might self-aggregating at concentration higher than 500nM (33). As for model membranes experiments, this self-aggregation will strongly impact our measurements and lead to kinetics reaction scheme totally different from the one we used to develop our numerical simulations and our empirical analytical solution (eq.6).

**Fig. 3.**
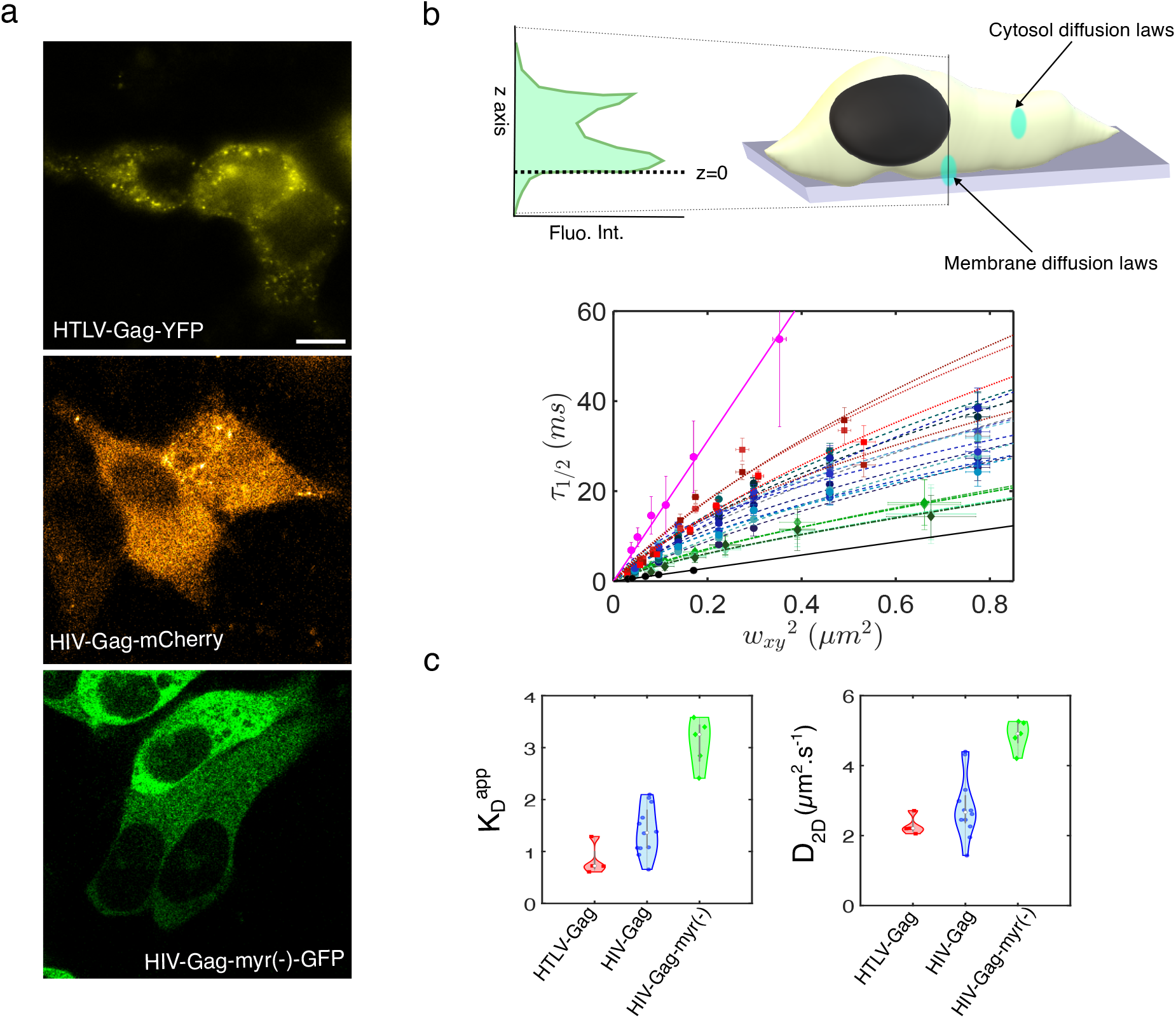
svFCS diffusion of different viral Gag proteins in HEK-293T cells. a- Typical confocal images obtained in HEK 293-T cells expressing, from top to bottom, HTLV-1-GagYFP, HIV-1-GagmCherry and HIV-1-myr(-)GagGFP from top to bottom. Scale bar is 10 *µ*m for the three images. b- Experimental diffusion laws obtained in cells expressing the different fluorescent viral Gag proteins. Red squares are HTLV-1-GagYFP data, blue circles are HIV-1GagmCherry data and green diamonds are HIV-1-myr(-)GagGFP data with their respective fits (dots, dashed, dot-dashed lines) using eq.6 with *K*_*D*_^app^ and *D*_2D_ as free parameters. Pink dots and line represent svFCS data and their fit using a linear model obtained at the plasma membrane of Bd-PC lipid labelled HEK-293T cells (n=2). Black dots and line represent svFCS data and their fit using a linear model of a cytosolic HTLV-1-GagYFP. c- Comparison of the *K*_*D*_^app^ and *D*_2D_ values obtained from the fit of the svFCS diffusion laws for the three different viral Gag expressed in HEK-293T cells.

Based on our ACF fits with eq.3 or eq.4, we found the average number of molecules (N) in the smallest waist to be between 3 and 20 which corresponds to apparent concentrations ranging between 35 to 230 nM (the average number of molecules found in the different confocal volumes are shown in supplemental figure SI.3a).

As we did for model membranes, we first fitted our svFCS data with eq.6 leaving the three parameters *K*_*D*_^app^, *D*_2D_ and *D*_3D_ free. As found previously, these fits systematically led to erroneous values of *D*_3D_ (*D*_3D_ = 10^4^ *µ*m^2^.s^*−*1^ in the case of HIV-1-Gag-mCherry). To circumvent that pitfall, we therefore measured *D*_3D_ directly into the cell, by performing svFCS with the laser focused in the cytosol, far from the plasma membrane (see black line and dots in figure 3b and SI.3 for examples). Linear fit of the *τ*_1*/*2_ values led to an estimated average *D*_3D_ value of 18.6*±*1.3 *µ*m^2^.s^*−*1^ (mean *±* sd, n=4 cells) *D*_3D_ was therefore fixed to 19 *µ*m^2^.s^*−*1^ and the data were fitted with *K*_*D*_^app^ and *D*_2D_ parameters left free. Figure 3b shows the experimental diffusion laws obtained in 4 cells expressing HTLV-1-GagYFP (redish squares), 12 cells expressing HIV-1-GagmCherry (blueish circles) and 5 cells expressing HIV-1-myr(-)GagGFP (greenish diamonds) with their respective fits as described above. As for model membranes, each of these diffusion laws are flanked by the protein cytosolic free diffusion laws (black dots and line) and by TopFluor®PC labelled plasma membrane (pink dots and line). In these plots, one sees three distinct sets of curves, corresponding to the three different proteins. These differences are confirmed when looking at the *K*_*D*_^app^ and *D*_2D_ values obtained from the fits (Fig. 3c).HIV-1-myr(-)Gag exhibits larger *K*_*D*_^app^ and *D*_2D_ values than the two other proteins, *K*_*D*_^app^=3.1*±*0.5 (mean *±* sd) and *D*_2D_=4.9*±*0.4 *µ*m^2^.s^*−*1^ (mean *±* sd). This implies that less than 20 % of the HIV-1-myr(-)Gag is bound to the plasma membrane of HEK293T cells. HIV-1-Gag and HTLV-1-Gag exhibit values in the same range. The membrane diffusion coefficients *D*_2D_ were found to be 2.2*±*0.3 *µ*m^2^.s^*−*1^ for HTLV-1 Gag and 2.6*±*0.9 *µ*m^2^.s^*−*1^ for HIV-1 Gag. These value are higher than the diffusion coefficient obtained from diffusion laws of TopFluor®PC (*D*_2D_=1.5*±*0.2 *µ*m^2^.s^*−*1^). The mean *K*_*D*_^app^ values were found to be 0.8*±*0.3 and 1.4*±*0.5 for HTLV-1 Gag and HIV-1 Gag respectively. Hence,on average, at this Gag concentrations 55*±*9% of HTLV-1 Gag molecules are found bound to the plasma membrane of HEK-293T cells, while it is only 40*±*14% in the case of HIV-1-Gag, revealing a higher affinity of HTLV-1-Gag for the plasma membrane.

## Discussion - Conclusion

Monitoring and quantifying molecular motions using FCS mainly rely on the ability to derive analytical solutions in order to fit the autocorrelograms obtained from single spot FCS measurements. Except in simple cases, such as flow or free diffusion, single point FCS often fails to correctly determine and quantify molecular motions in heterogeneous and non-isotropic environment. This failure is frequently (and sometimes abusively) circumvented by the addition of an *α* exponent, signing for anomalous sub-diffusion motion in the heterogeneous media. However, anomalous sub-diffusion may occur from many different processes which theoretically cannot be uniquely identified or quantified by the sole value of *α* (see (34, 35) and references therein). In the case of reaction-diffusion dynamics, it has been shown that no simple analytical solutions could be derived to fit single spot FCS experiments (3). This is also what we found here. Deriving an expression systematically required to simplify the dynamics process with different hypotheses, as was also observed in (3). In the 2D/3D diffusion and binding dynamics, we expected to have two limit regimes, namely pure 3D free diffusion (membrane unbound molecule, *k*_off_ ≫ *k*_on_) and pure 2D diffusion (membrane bound molecule, *k*_off_ ≪ *k*_on_) as well as different intermediate regimes where the system dynamically equilibrates. Interestingly, the fit of the autocorrelograms with different analytical expressions (3D free diffusion, 2D free diffusion, 2-components diffusion and anomalous diffusion) shown that this type of dynamics is hardly correctly fitted by anomalous diffusion if, in order to select the best-fitting expression (as is the case with information-theory derived criteria), the number of free parameters to fit is accounted for. This is in line with experimental data obtained on a membrane binding exchange factor PH domain probed by FRAP experiments, which dynamics was also shown not to be anomalous (36).

A powerful method to correctly probe and characterize the dynamics of molecules in complex media is the spot size variation method (5, 11) (sv-FCS) that establishes FCS diffusion laws. We therefore used computer simulations to generate synthetic FCS diffusion laws for 2D/3D diffusion + binding in different regimes. We first controlled that these simulated diffusion laws were flanked by the two expected limit regimes, namely 2D and 3D diffusion, that both lead to linear FCS diffusion laws. We then generated a set of synthetic diffusion laws in the intermediate regime, by varying the values of *D*_2D_, *D*_3D_ and *K*_*D*_ in our simulations. In these intermediates regimes, we systematically obtained non linear synthetic FCS diffusion laws. We therefore derived an empirical non-linear analytical expression (eq.6) that, when fitted to our synthetic diffusion laws, provides a a quantitative estimation of *K*_*D*_ and *D*_2D_, with respective relative precision of 24% and 14%. While we agree that this precision can be considered as moderate, we stress out that obtaining the absolute value of a parameter in biology is not always a more important information than correctly estimating its variation with changing biological conditions.

For this reason, we challenged the capacity of our empirical expression to estimate the change of the membrane apparent binding affinity *K*_*D*_^app^ and the membrane diffusion *D*_2D_ occurring when HIV-1 and HTLV-1 Gag proteins and a HIV-1 derivative (myr(-)) binds either to model lipid membranes, with controlled composition, or at the plasma membrane of living HEK-293T cells. In both cases, fitting the experimental FCS diffusion law with (eq.6) did not provide with correct estimates when we tried determine the three parameters of the model simultaneously, namely *D*_3D_, *D*_2D_ and *K*_*D*_. The simplest explanation, is that the number of beam waists to fit is not sufficient to achieve a good estimation of the three parameters at the same time. Increasing the number of waists monitored should help in having better fits, but while it is easy to achieve numerically, it remains illusive experimentally. Indeed, correct determination of *τ*_1*/*2_ needs avoiding photo-bleaching during successive experiments, which cannot be done if we drastically increase the number of waists probed in the same cell. To circumvent that issue, we directly measured the *D*_3D_ in the bulk (in the case of model membranes) or in the cytosol (in the case of HEK293T cells). Once *D*_3D_ determined, we could successfully fit the experimental svFCS diffusion laws the two free parameters remaining (*K*_*D*_^app^ and *D*_2D_). We first used SLBs model membranes made of PC, PS and PI(4,5)P_2_ with decreasing surface negative charges by tuning the molar ratio of PS and PI(4,5)P_2_. We found the membrane diffusion *D*_2D_ of HIV-1-myr-Gag to be equivalent to that of Atto-647N-PI(4,5)P_2_ in the SLB. PI(4,5)P_2_ is known to be the specific target lipid of HIV-1-Gag association to the plasma membrane (31) and has been shown to be trapped by HIV-1-Gag in model membranes (32) as well as in living T-cells (37). However, PS is also involved in binding of Gag to the membrane (18, 19). This could explain why we only measured *K*_*D*_^app^>1 for SLBs lacking negatively charged lipids. Although surprising, as we expected to obtain increasing values of *K*_*D*_^app^ with decreasing amount of PI(4,5)P_2_, the *K*_*D*_^app^ values obtained with svFCS diffusion laws exhibited exactly the same trend than the those obtained by LUV binding experiments (i.e., *K*_*D*_^app^>1 only observed with neutral lipids). We also previously showed that lack of myristate strongly decreases the specificity for PI(4,5)P_2_ (19). In addition, in a 2:1 PC:PS mol:mol lipid composition, (38) reported no significant change in membrane binding for molar proportion of PI(4,5)P_2_ varying from 0 to 2%, in agreement with our own measurements.

Finally, we examined the ability of our method to quantify membrane binding and diffusion of retroviral Gag proteins and its derivatives in HEK-293T cells. Interestingly, we found *D*_2D_ to be 2.3*±*0.3 *µ*m^2^.s^*−*1^ and 2.8*±*0.8 *µ*m^2^.s^*−*1^ for HTLV-1 and HIV-1 Gag, respectively, which is in the range of the bound *D*_2D_ found on model membranes and in the range of the values obtained in the same cells for TopFluor®-PC, a neutral membrane lipid. This confirms that our method led to a correct estimation of the membrane bound diffusion coefficient of retroviral Gag proteins even in living cells. Using a combination of TIRF and fluorescence fluctuation spectroscopy, (21) showed that HTLV-1-Gag has a higher affinity than HIV-1-Gag for the plasma membrane of HeLa cells. Using svFCS diffusion laws, we also measured a lower *K*_*D*_^app^ for HTLV-1-Gag compared to HIV-1-Gag. This reflects the higher affinity of HTLV-1-Gag vs HIV-1-Gag for the plasma membrane of HEK-293T cells and illustrates again the ability of our method to correctly determine this parameter. We also showed that removing myristate from HIV-1-Gag resulted in the re-localization of the Gag protein towards the cytosol, with only 24*±*3% of the total HIV-1-myr(-)Gag proteins bound to the membrane. Again this value is in good agreement with the results previously obtained by (21). Importantly, the results we obtained here are in perfect line with previous results obtained by different groups using different approaches/techniques.

It is not the main aim of this study to establish a molecular mechanism for retroviral Gag proteins by accurately quantifying their binding to lipid membranes in different conditions. However, we believe it is worth stressing out that not only we have strong differences in the *K*_*D*_^app^ values obtained for HIV-1-myr(-)Gag in model vs cells lipid membranes but also a lower *K*_*D*_^app^ in the case of HIV-1-Gag in cells vs. HIV-1-myr(-)Gag on negatively charged SLBs. Several reasons could account for such discrepancies between model lipid membranes and living cell plasma membranes :

- The accessibility of PI(4,5)P_2_ in the plasma membrane of the cells, and more generally of negatively charged lipids, might be strongly decreased compared to model membranes, as many other proteins present in the cells are also known to interact with these lipids. This would result in a screening of the targeted lipids and, consequently increase *K*_*D*_^app^ values as *c* decrease.
- In the presence of RNA (which are highly present in cells), HIV-1 Gag has been shown to adopt a horseshoe configuration where both MA and NC domain binds to the RNA (39). The membrane binding process is then mediated by the HBR domain of the MA interaction with PI(4,5)P_2_, inducing the release of the RNA which stays bound to the NC domain (40). This screening of HIV-1-Gag membrane binding domain in cells might be more important than in our model membrane experiments, where RNA is hardly present, resulting again in an increase of *K*_*D*_^app^.
- In the absence of RNA, the NC domain has been shown to exhibit significant affinity for negatively charged lipids (41). In our model membranes, where lipids are in large excess compared to RNA, this second lipid binding motif in the NC domain of HIV-1-Gag could compensate the lack of myristate and favor stronger binding to the SLB (lower *K*_*D*_^app^), while it will be screened in cells by the cytosolic/cellular RNAs.

In this study, we have demonstrated the ability of svFCS diffusion laws to better estimate (apparent) membrane binding coefficients and membrane diffusion coefficients than single spot FCS can. Using a numerical simulation-based approach, we have derived an empirical analytical expression that we then used to fit experimental svFCS data obtained on retroviral Gag proteins binding either to model membranes or to plasma membrane of living cells. Overall, the results we obtained with our method are in perfect line with the ones reported in previous literature, obtained with different methods. We conclude that our results provide a non-invasive and direct way to fairly estimate, in the same experiment, membrane binding and diffusion coefficients in living cells.

## Author Contributions

CF, HB & DM designed the project and the experiments. HB developed the computer code, performed the numerical simulations and derived the empirical expression. AM, EB and JN performed the FCS experiments. AM, EB and JN fitted experimental data. RD & CA prepared the cell samples, EB the SLBs. JM purified and provided the myr(-)Gag protein. HB & CF draft the manuscript assisted by AM, EB, CA & DM.

## Acknowledgments

The authors acknowledge the Imabio CNRS (GdR Imabio) consortium for their continuous support of the project and for inititally granting EB. AM Ph.D. fellowship is granted by CNRS Prime 80. EB was then granted by CNRS. CA is a recipient of Université Montpellier Ph.D. fellowship. RD Ph.D. fellowship is granted by Sidaction. The project was initially granted by ANR Fluobuds and then by CNRS. Authors acknowledge Montpellier RIO Imaging (MRI, Biocampus, UAR CNRS) microscopy facility. Authors acknowledge Eggeling, H.de Rocquigny, M. May, D. Derse and N. Jouvenet for the gift of the different fluorescent lipids, dyes and plasmids.

## Supporting Information

### SI1. Derivation of the auto-correlation function for the single-spot case

#### Main Hypotheses

Here we simplify the problem by considering that the membrane binding and diffusion sites are not restricted to at *z* = 0 but are homogeneously distributed in the 3D bulk. Moreover, we neglect the reduction of dimensionality and consider that the bound molecules diffuse in the same 3D space as the free ones, but with a different diffusion coefficient. Under these assumptions, the problem is now that of a protein that reversibly switches between two forms: (F)ree and (B)ound, and where the two forms have non-identical but non-zero diffusion coefficients: 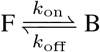, with *D*_3 *D*_ > *D*_*D* 2_ > 0.

Assuming Brownian motion in 3D (for *both free and membrane-bound* Gag), yields the following reaction diffusion system:

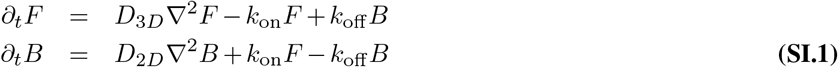

where *F* and *B* refer to the local concentration of the two species.

#### General case

Noting *f* and *b* the fluctuations of *F* and *B*, respectively (i.e. 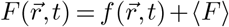 and 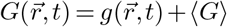 where ⟨*F*⟩ and ⟨*G*⟩ are the average concentrations over the volume), Eq. SI.1 yields:

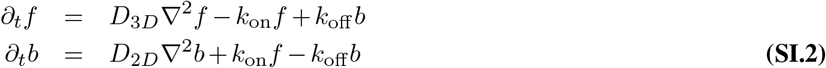

The coefficient matrix of the Fourier transform of Eq. SI.2, *M*_*ij*_ = *K*_*ij*_ *− D*_*i*_*q*^2^*δ*_*ij*_ (where *K* is the stoechiometry matrix and *q*^2^ = *q*_*x*_^2^ + *q*_*y*_^2^ + *q*_*z*_^2^ is the square norm of the Fourier variable) is given by:

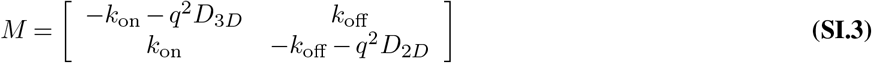

whose eigenvalues are:

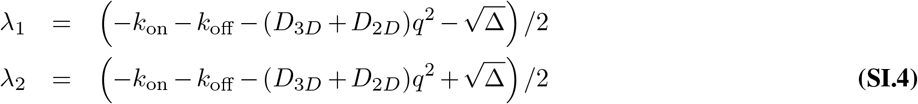

with Δ = *q*^4^(*D*_3*D*_ −*D*_2*D*_)^2^+(*k*_on_+*k*_off_)^2^+2*q*^2^(*D*_3*D*_ −*D*_2*D*_)(*k*_on_−*k*_off_).

The corresponding eigenvectors are

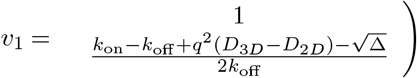

and

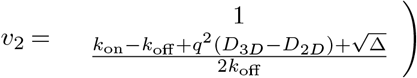

Their inverse eigenvectors are

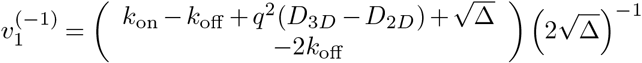

and

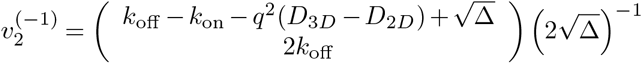

Assuming a Gaussian point-spread function:

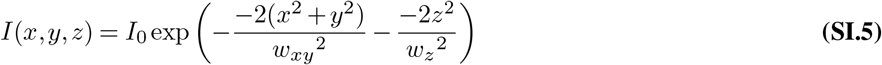

the autocorrelation of the fluctuations *G*(*τ*) can be generically computed (27) as :

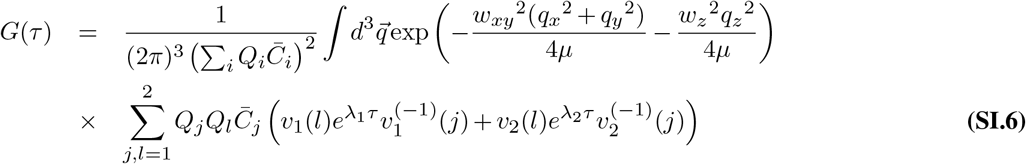

where *µ* = 1 or 2 for one- or two-photon fluorescence, respectively, *Q*_*i*_, *i* = 1, 2 are the fluorescence cross-sections of F and B, respectively, 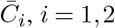 are their average distribution in the sample, i.e. 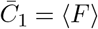 and 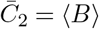, and *u*(*i*) represents the *i*th element of vector *u*.

#### Simplifications

##### Fast free diffusion

Eq.(SI.6) cannot be solved analytically without further simplifications. In particular, one can assume that the diffusion of F in the bulk is too fast to contribute a signal in FCS. In this case *Q*_1_ = *Q*_*F*_ *≈* 0 and the only non-vanishing terms in the sum of the rhs of eq.(SI.6) are for *j* = *l* = 2. Therefore in the specific case studied here, eq.(SI.6) becomes

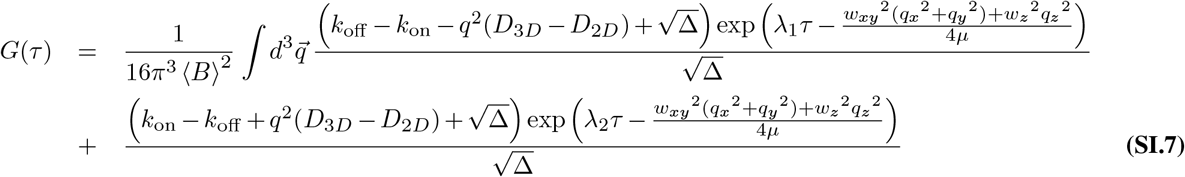

where the eigenvalues *λ*_*i*_ are given by eq.(SI.4). Eq.(SI.7) greatly simplifies when e.g. *D*_3*D*_ = *D*_2*D*_ since in this case, 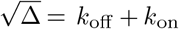 so that all the integral variables *q*_*i*_ are found inside the exponentials. The integral eq.(SI.7) can be analytically computed in this case and leads to the formula given in (27). In our case *D*_3*D*_ > *D*_2*D*_ > 0, though, we have not been able to compute the integral analytically.

In (3), the authors explore a series of parameter regimes where the expression for *G*(*τ*) can be further simplified.

##### The Hybrid Regime

This corresponds to the case where binding is much larger than unbinding, *k*_on_ *≫ k*_off_. In particular, this can be used to support the approximation *k*_on_ *− k*_off_ *≈ k*_on_ + *k*_off_. In this case, Δ simplifies according to

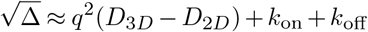

and

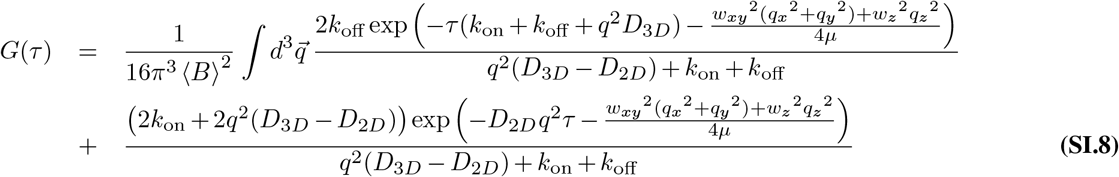

Unfortunately, this simpler expression is still too complex to permit the calculation of an analytical expression. *Other Regimes* None of the other regimes explored in (3) are applicable when *D*_2*D*_ > 0.

- The “pure diffusion” regime (*k*_on_ *≪ k*_off_) implies that all the Gag molecules are free. The case however is trivial, and corresponds to classical 3D diffusion in the bulk.
- The “effective diffusion” regime (*τ*_*D*_ = *w*_*xy*_^2^*/*(4*µD*) *≫* 1*/k*_on_) assumes that diffusion is slow enough that the binding reaction is at equilibrium (locally) everywhere and allows to reduce the reaction-diffusion system eq.(SI.2) to a single diffusion equation (no coupling to reaction). This reduction is however not possible when *D*_2*D*_ > 0
- the “reaction dominant” regime (*τ*_*D*_ *≪* 1*/k*_on_) assumes that the characteristic time for a Gag molecule to diffuse across the focal volume is much shorter than the characteristic time to bind the membrane. Here again, the simplification is critically based on the fact that one of the species does not diffuse and cannot be applied in our case.

### SI2. Supporting Figure

**Fig. SI.1.**
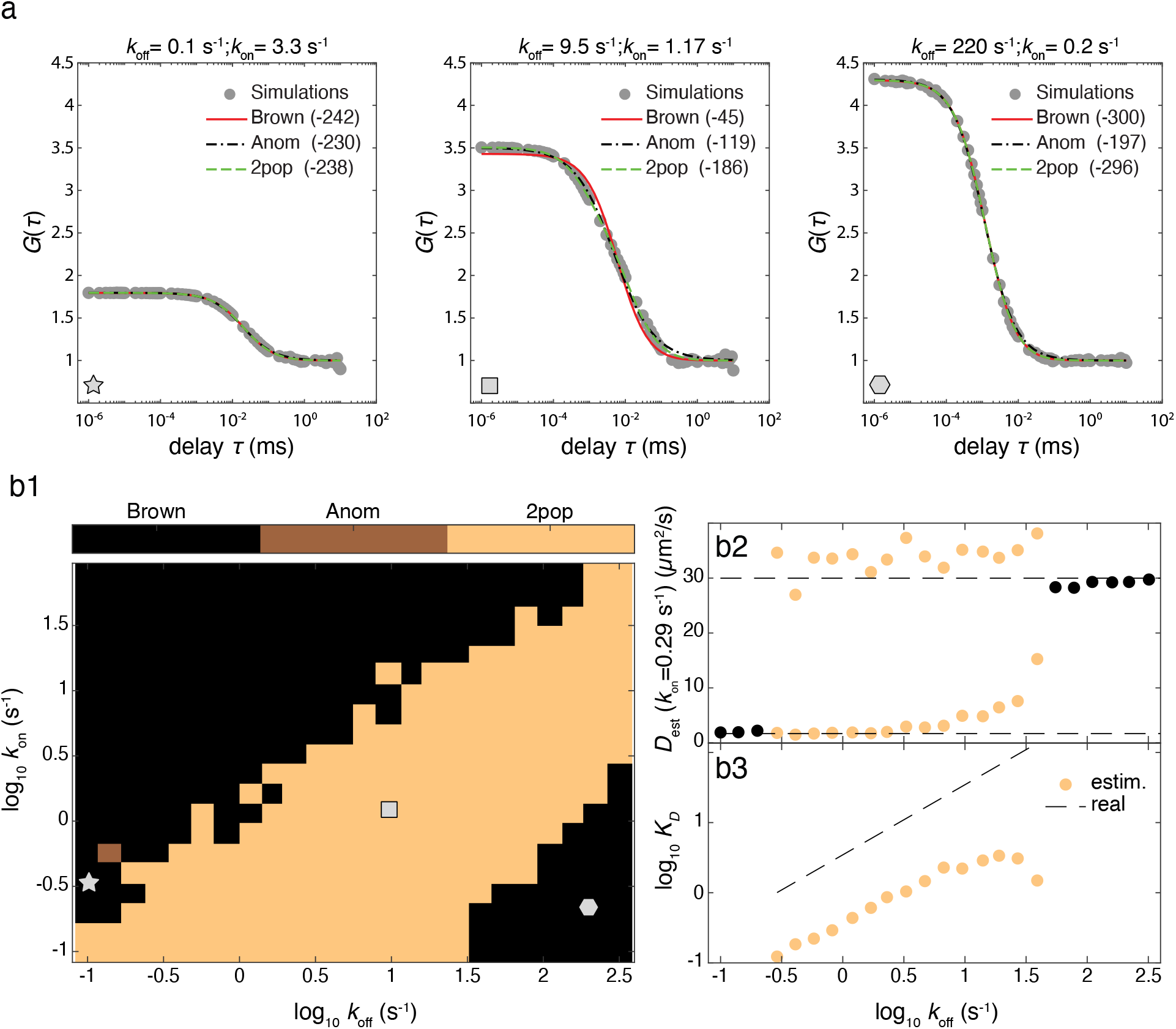
Fitting the auto-correlation function with adhoc expressions. Molecular diffusion in the cytoplasm and binding to the plasma membrane was simulated as described in Materials and Methods for a range of *k*_off_ and *k*_on_ values. We fixed the other parameters to constant values, in particular *w*_*xy*_ = 0.379 *µ*m, *D*_2*D*_ = 1.7 and *D*_3*D*_ = 30 *µ*m^2^*/*s. Illustration of some of the obtained auto-correlations are shown in (a) (full gray circles). Every auto-correlation functions were fitted by three models: a single population of pure 3D Brownian motion (full red line), a mix of two populations with pure 3D Brownian motion (dashed green line) or anomalous diffusion (dashed-dotted black line). The best model was selected by comparison of the corrected Akaike information criterias (28) (AICc, values shown in parenthesis). When comparing two models, the model with the smaller AICc is the best, with an evidence considered strong enough if the AICc difference was at least 6. In case several models were found equally good, we selected the model with the smaller number of free parameters. The colormap (b1) shows what is the best model for the range of *k*_off_ and *k*_on_ values explored. The gray star, rectangle and hexagon locate the illustrations shown in (a). With *k*_on_ fixed to 0.29 s^*−*1^, we also show the resulting estimation by the best model of the diffusion coefficient(s) (b2) and equilibrium constant *K*_*D*_ = *k*_off_ */k*_on_ (b3). The real values of diffusion and the equilibrium constants are shown as dashed lines.

**Fig. SI.2.**
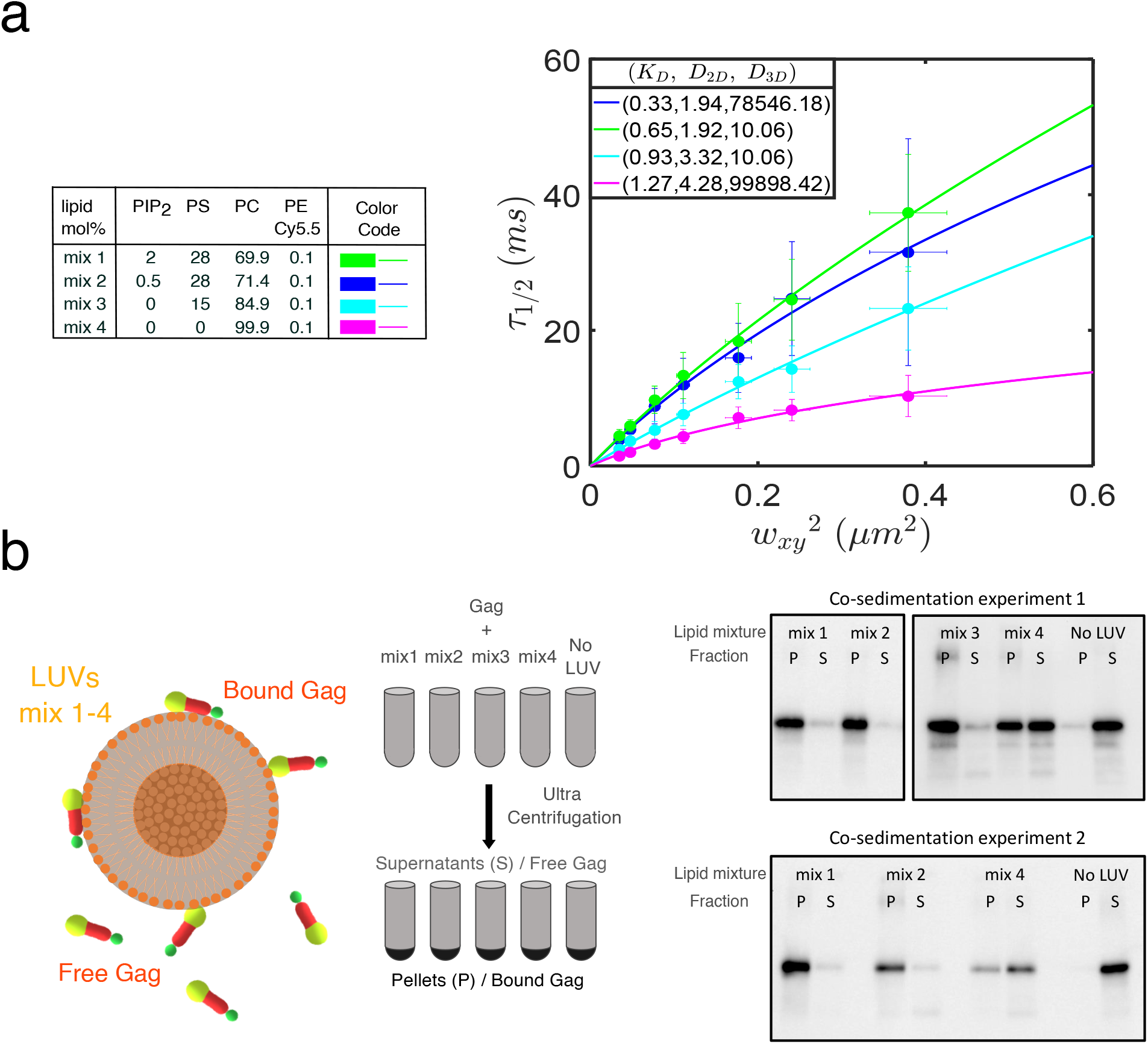
Supplemental results for Gag binding to model membranes. a- Fit of the FCS diffusion obtained with the different lipid mix (see table left for lipid composition) using a 3 parameters (*K*_*D*_^app^, *D*_2D_, *D*_3D_) free model. Although *K*_*D*_^app^ show increasing values with decreasing negative surface charge and PI(4,5)P^2^, the *D*_3D_ values obtained were totally incoherent. b- Principle of the LUV binding experiment (left) and images of the 2 western blots obtained in two different experiments and used to quantify the *K*_*D*_^app^ (see methods for details)

**Fig. SI.3.**
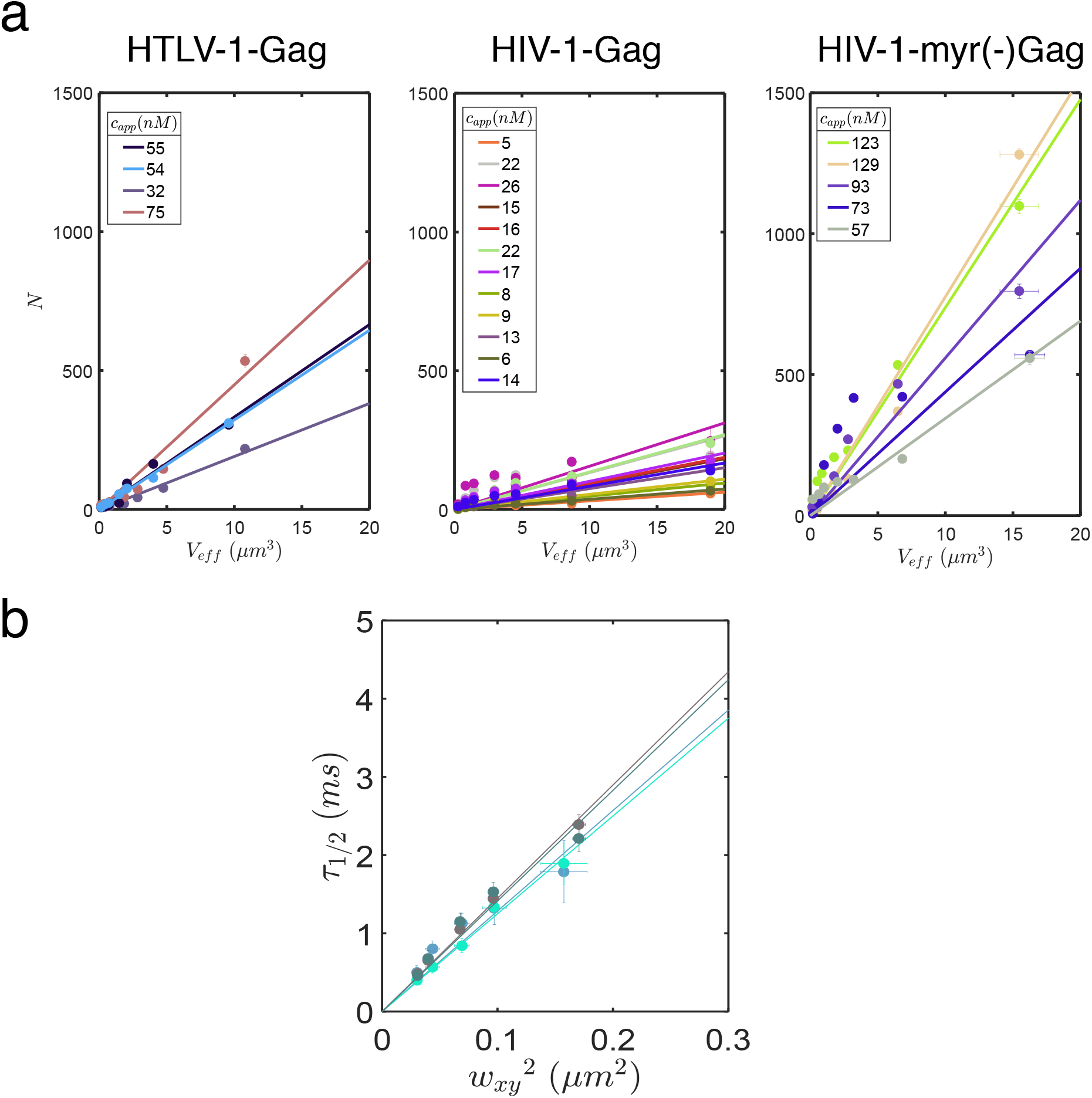
Supplemental results for Gag binding to cell plasma membranes. a- Evolution of the apparent number of molecules (N) obtained from the fit of ACF with eq.3 or eq. 4 (see methods), as a function of the effective volume (*V*_*eff*_) for all the cells analyzed here. The slope of each curve is the apparent concentration constantly increasing from HIV-1-Gag-mCherry to HIV-1-myr(-)Gag-YFP, while still being (far) below the threshold concentration of self-assembly (see text for detailed explanations). This apparent concentration is systematically found lower than the one estimated at the smallest waist. This is mainly due to the fact that at big waists, the apparent number of molecules N growing, it is difficult to be correctly estimated and leads to underestimation of apparent concentrations. b- Example of FCS diffusion laws obtained in the cytosol of 4 different cells, in the case of HTLV-1-Gag-YFP (shades of grey dots) and HIV-1-myr(-)-Gag-YFP (shades of cyan dots) and their linear fit (lines of same colors) assuming 3D free diffusion. Slopes of the fits are used to determine *D*_3D_.

